# Gametogenesis in the green seaweed *Ulva mutabilis* coincides with massive transcriptional restructuring

**DOI:** 10.1101/2021.08.13.456063

**Authors:** Xiaojie Liu, Jonas Blomme, Kenny Bogaert, Sofie D’hondt, Thomas Wichard, Olivier De Clerck

**Author notes:** Email addresses: Xiaojie Liu, Jonas Blomme, Kenny Bogaert, Sofie D’hondt, Thomas Wichard. Olivier De Clerck (Corresponding author).

## Abstract

The molecular mechanism underlying sexual reproduction in land plants is well understood in model plants and is a target for crop improvement. However, unlike land plants, the genetic basis involved in triggering reproduction and gamete formation remains elusive in most seaweeds, which are increasingly viewed as an alternative source of functional food and feedstock for energy applications. Here, gametogenesis of *Ulva mutabilis*, a model organism for green seaweeds, is studied. We analyze transcriptome dynamics at different time points during gametogenesis following induction of reproduction by fragmentation and removal of sporulation inhibitors. Analyses demonstrate that 45% of the genes in the genome are differentially expressed during gametogenesis. We identified several transcription factors that potentially play a key role in the early gametogenesis of *Ulva* given the function of their homologs in higher plants and microalgae. In particular, the detailed expression pattern of an evolutionary conserved transcription factor containing an RWP-RK domain suggests a key role during *Ulva* gametogenesis. The identification of putative master regulators of gametogenesis provides a starting point for further functional characterization.

**Highlight:** Transcriptomic analyses of gametogenesis in the green seaweed *Ulva* highlight the importance of a conserved RWP-RK transcription factor in induction of sexual reproduction.

## Introduction

Sexual reproduction is one of the most important and conserved processes in eukaryotes. In essence, sexual reproduction encompasses the fusion of two haploid gametes of opposite sex to form a diploid zygote. Zygote formation is either followed by meiosis to restore the haploid state (zygotic meiosis) or by development of a diploid life stage through mitotic divisions (gametic meiosis) (Coelho *et al.*, 2007). Molecular genetic studies of reproduction in land plants resulted in an excellent understanding of the fundamental biological principles and genes involved. However, unlike land plants, the molecular mechanisms involved in the onset of reproduction, the formation of gametes (gametogenesis), fusion and meiosis remain elusive for most algae and seaweeds. In the green algal lineage, the molecular mechanisms underlying gametogenesis and fusion have been extensively studied in *Chlamydomonas reinhardtii*, a unicellular freshwater green alga (Abe *et al.*, 2004; Abe *et al.*, 2005; Goodenough *et al.*, 2007; Lin and Goodenough, 2007), but to which extent key regulatory genes are conserved across green algae or even land plants remains unclear.

Here, we study the expression of genes during the various stages of gamete formation in *Ulva*, an emerging model for green seaweeds (Balar and Mantri, 2020; De Clerck *et al.*, 2018; Wichard *et al.*, 2015). *Ulva* species are abundant in coastal benthic communities around the world and form a potential source of biomass that can be used for food, feed, biofuel and nutraceuticals (Alsufyani *et al.*, 2014; Balar and Mantri, 2020; Bolton *et al.*, 2016; Neori *et al.*, 2004; Nisizawa *et al.*, 1987; Wang *et al.*, 2019). Given its commercial value, reproducibly controlling the life cycle of *Ulva* is important in the development of this crop. Under natural conditions, large parts of the *Ulva* thallus are converted into gametangia or sporangia in mature individuals and the entire cell content is converted into reproductive cells. Numerous studies have investigated the effect of different endogenous (e.g. production of sporulation and swarming inhibitors) and environmental factors (e.g. light, temperature or nutrient conditions) on gametogenesis in different *Ulva* species (Luning *et al.*, 2008; Mantri *et al.*, 2011; Sousa *et al.*, 2007; Stratmann *et al.*, 1996; Wichard and Oertel, 2010). Although we are still far removed from a complete understanding of the interplay of the various factors that appear to play a role, thallus fragmentation has been recognized as the most efficient way to induce gametogenesis (Gao *et al.*, 2010; Hiraoka and Enomoto, 1998; Stratmann *et al.*, 1996; Vesty *et al.*, 2015; Zhang *et al.*, 2011).

The ability to efficiently induce gametogenesis in laboratory cultures of *Ulva mutabilis* provides an elegant time course to study both phenotypic and molecular factors involved in this process (Wichard and Oertel, 2010). Two sporulation inhibitors, SI-1 and SI-2 control the onset of gametogenesis of *Ulva mutabilis.* SI-1 is a species-specific glycoprotein secreted in the culture medium which depletes with maturation of the thallus. SI-2 is a nonprotein molecule existing within the inner space between the two blade cell layers. Thallus fragmentation followed by a washing step whereby spent culture medium is replaced, results in the total removal of sporulation inhibitors and induces gametogenesis (Stratmann *et al.*, 1996). Following fragmentation and washing, SI-1 and SI-2 are removed and vegetative cells enter a determination phase (~0-26h, Fig. 1) in which a ‘swarming inhibitor’ (SWI) is produced (Wichard and Oertel, 2010). SWI functions as a mechanism to synchronize gamete release. The start of the differentiation phase coincides with the completion of the S-phase and the G2-phase (~36 h) in the first cell cycle (Fig. 1). At the end of the determination phase, the cells enter the next S-phase and become irreversibly committed to gametangium differentiation. The differentiation phase (~26-70h) is characterized by a reorientation of the chloroplast, followed by four consecutive cell divisions forming sixteen progametes per cell and their maturation. Mature gametes are eventually released during the swarming phase, following a light stimulus or if the SWI declines in concentration on the third day after induction (Kessler *et al.*, 2017; Wichard and Oertel, 2010). Given that the whole process of *Ulva* gametogenesis, can be subdivided in discrete phases, the gene regulatory networks involved in the different stages of gametogenesis should be distinct. As a first step toward understanding the molecular mechanisms of gametogenesis, we established the transcriptional structure for *Ulva mutabilis* gametogenesis in this research. We make use of differential removal of SI’s to separate the effect of fragmentation on transcription from gametogenesis. Transcription patterns are compared with those of other green algae and land plants.

**Figure 1.**
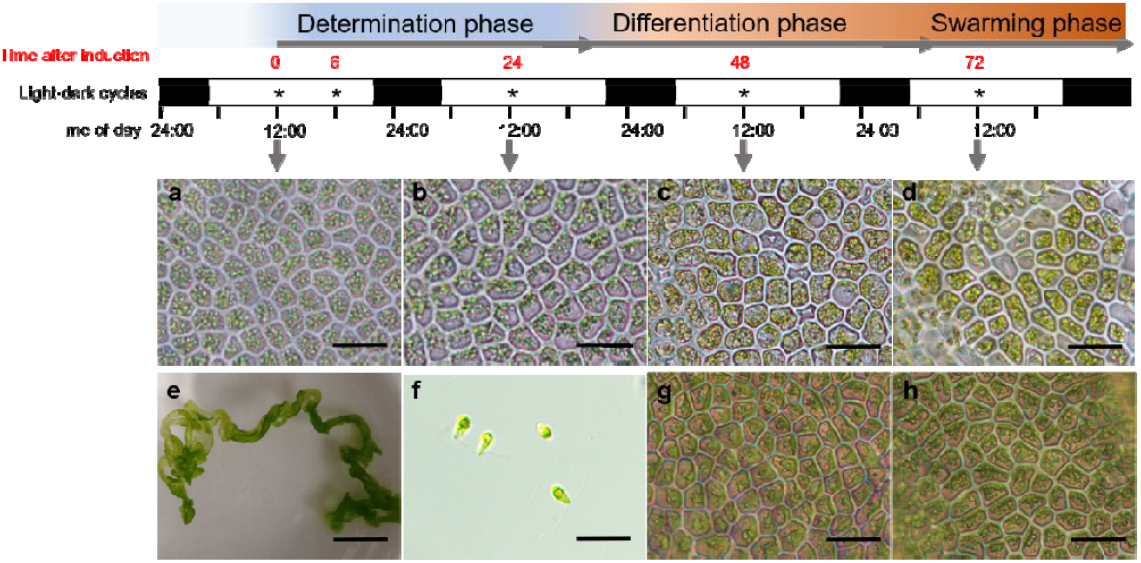
Time course of gametogenesis in *Ulva mutabilis.* “determination phase” and “differentiation phase” described previously by Stratmann et al. (1996). The “swarming phase” is the time when the change of medium can release the gametes. Asterisks indicate the sampled time points for RNA-seq. (a) vegetative cells before induction, (b) gametogenesis induction after 24 h with chloroplast reorientation, (c) gametangia after 48 h, (d) reproductive cells with fully formed gametes and discharged gametes after 72 h, (e) *Ulva* thallus before induction, (f) released gametes, (g,h) induced *Ulva* tissue without refreshment of growth medium after 48 (g) and 72 hours (h), demonstrating that no gametes were formed as long as sporulation inhibitors are not removed. Scale bars: 2cm (e) or 20 μm (others).

## Materials and methods

### Strains and Culture Conditions

Haploid gametophytes of *Ulva mutabilis* [strain ‘wildtype’ (wt-)] were collected initially in southern Portugal (Føyn, 1958). Gametophytes of *U. mutabilis* were raised parthenogenetically from unmated gametes and cultured in transparent plastic boxes containing 2 L Provasoli enriched seawater (PES) medium (Provasoli, 1968) with the following specific parameters: light intensity, 70 μmol photons m^-2^·s^-1^; temperature, 18 ± 1 °C; and 17:7 h light:dark cycle. The medium was completely changed every 2 weeks until fertility (~10 weeks). Afterwards, the medium was partially (20 %) changed to avoid induction of gametogenesis.

### Induction of gametogenesis

Intact mature thalli of *U. mutabilis* were cut into 1-3 mm^2^ fragments using a herb chopper. Fragmentation was carried out in the morning. Fragments derived from a single individual were separated into 2 groups. Fragments were washed 3 times for 15 minutes with autoclaved seawater and transferred to new medium (CW, chopping and washing treatment). The washing step removes sporulation inhibitors thereby inducing differentiation of the gametangia (Stratmann *et al.*, 1996; Wichard and Oertel, 2010). Triplicates of 0.2 g fragmented thallus were collected at 0, 6, 24, 48 and 72 h following gametogenesis induction. Control treatments consisted of thalli that were fragmented but not washed and transferred immediately to the original culture medium (CNW, chopping no washing). The control was sampled at 6 and 24 h, respectively. The presence of SI-I inhibits gametogenesis and control thalli therefore remain ‘vegetative’ after 72 h. A third and a fourth treatment, used for expression analyses of RWP-RK transcription factors, consisted of thalli with normal growth that were neither chopped nor washed (NCNW) or not chopped but washed (NCW), respectively. Samples were flash-frozen in liquid nitrogen and stored at −80 °C for RNA extraction.

### RNA extraction and RNAseq

Total RNA was extracted using a CTAB method (Le Bail *et al.*, 2008). Quality and quantity of total RNA was evaluated using a NanoDrop™ 2000c spectrophotometer (Thermo Fisher Scientific) and Bioanalyzer RNA6000 (Agilent Technologies). cDNA libraries were constructed with a Quantseq™ 3’ mRNA-Seq library prep kit (Lexogen) following the manufacturer’s instructions. Illumina sequencing was performed using the Illumina Nextseq 500 platform to produce 50 bp single-end reads. RNA reads are available as SRA-xxx. Reads were mapped to the *U. mutabilis* genome (De Clerck *et al.*, 2018). We used the version of the *slender* strain which has UTR-regions annotated, as opposed to the *wildtype* genome. Reads were mapped using TopHat ver. 2.1.1 (Trapnell and Salzberg, 2009) with default parameters. The number of mapped reads was calculated using HTseq (Anders *et al.*, 2015). Differentially expressed genes (DEG) were identified between each treatment and control groups using the R package *edgeR* (Robinson *et al.*, 2010; Team, 2013). The sample variation was estimated by tag-wise dispersion. Raw counts were normalized by CPM (counts per million reads mapped). Genes whose CPM value was >1 were considered as ‘expressed’. A false discovery rate (FDR) of 0.05 and absolute value of log 2-fold change of > 1 were adopted as thresholds for differential expressed genes (DEGs) detection.

### Hierarchical Clustering Analysis

To study the transcriptional dynamics during the gametogenesis process, DEGs between any two of five time points (0 h, 6 h, 24 h, 48 h, 72 h) of the treatment group were identified. The identified DEGs were subjected to a hierarchical clustering analysis by Pearson correlation (Corpet, 1988) based on their CPM values in the R (ver. 3.5.0) programming environment using the *hclust* function. Prior to the analysis, the optimal number of clusters was identified and investigated by performing a SSE merit analysis and an R-based average Silhouette Width analysis (Langfelder *et al.*, 2008). SSE is defined as the sum of the squared distance between each cluster member and its centroid cluster. As the number of clusters increases, the distance between each point and its centroid will be smaller. The optimal number of clusters is suggested when the addition of a new cluster does not significantly decrease the SSE. The silhouette value describes how similar a gene is to its own cluster (cohesion) compared to other clusters (separation).

### Functional Annotation and GO enrichment

Transcription factor (TF) identification and GO terms for gene models were retrieved from the *Ulva mutabilis* genome annotation file (downloaded from https://bioinformatics.psb.ugent.be/orcae/). The top hit of homologs acquired by the integrative orthology method of the differential expressed TFs of *A. thaliana* and *C. reinhardtii* were obtained using a custom-built PLAZA-*Ulva* version (Vandepoele *et al.*, 2013). The functional annotation of these homologs was acquired on TAIR (https://www.arabidopsis.org/) or Phytozome (https://phytozome.jgi.doe.gov/pz/portal.html). GO enrichment analysis for DEGs was based on a Fisher’s exact test implemented in the *TopGO* package in R (Alexa and Rahnenführer, 2009). The enriched GO terms (p<0.01) in the category “biological function” were summarized using the REVIGO web server, which performed a clustering algorithm that relies on the semantic similarity method (Supek *et al.*, 2011).

### TF identification analysis

To verify RNA-seq results and to obtain more detailed expression pattern for specific TFs during gametogenesis, we performed RT-qPCR analysis of the expression of up-regulated TFs. We sampled the treatment group (CW, described above) and thalli with normal growth that were neither chopped nor washed (NCNW) every 2 h in the first 48 h during gametogenesis to obtain detailed expression profiles for TFs identified in the initial RNA-seq experiment. We designed an experiment consisting of four groups: normal vegetative thalli without chopping nor washing (NCNW); chopped thalli that were not washed (CNW); chopped and washed thalli (CW, normal induction); intact thalli that were washed to remove of SI in the medium (NCW). Gametogenesis was partly induced in the latter treatment. Thalli were sampled at 6 h for the four treatments to identify the expression profile of the TFs. RNA was extracted as described above and cDNA was synthesized using an iScript™ cDNA Synthesis Kit (Bio-Rad). PCR was performed using Bio-Rad CFX96 Real-Time PCR systems. Reactions were performed in a final volume of 10 μL containing 5 μL of SYBR Green Master Mix, 0.5 μM each primer, and 10 ng of cDNA under following program: 5 min at 95 °C followed by 35 cycles of 95 °C for 10 s and 55 °C for 30 s. Gene expression levels were calculated using the ddCt method (Livak and Schmittgen, 2001). *UBIQUITIN* (*UMSL012_0173*) and *ELFA* (*UMSL 016_0119*) were stable under a wide range of time points and were taken as reference. Primer sequences are listed in Table S1.

## Results

To dissect the molecular mechanisms underlying *Ulva* gametogenesis, the global gene expression levels of *U. mutabilis* were measured by RNA-Seq as a function of time. We sampled five-time points: 0 h, 6 h, 24 h, 48 h and 72 h after induction of gametogenesis (Fig. 1) under standard induction conditions (CW) and two-time points (6h and 24h after induction of gametogenesis) that omitted the washing step (CNW), with three biological replicates for each time point and condition. RNA-sequencing resulted in 215 x 10^6^ reads, >70% mapped to the *U. mutabilis* genome. The total number of mapped reads for each sample and the raw reads counts for each gene were listed in Table S2 and Table S3, respectively. Transcript profiles were highly reproducible among the three biological replicates at each time point based on the Pearson R^2^ test and PCA analysis (Table S4, Fig. S1).

### Gene expression pattern during gametogenesis

In total, 8296 distinct genes (62.2% of annotated genes) were expressed during gametogenesis, with relatively low variation in the total number of expressed genes between the time points: ranging between 7146 (0 h) and 7949 expressed genes (72 h). We identified 6056 differentially expressed genes (DEGs) between any two of the five time points (0 h, 6 h, 24 h, 48 h, 72 h) during the gametogenesis process, representing 45% of genes of the annotated genome. Each time point could be characterized by a set of genes describing the various steps of gametogenesis (Fig. 1). Hierarchical clustering grouped genes in 5 clusters for further analyses based on the stabilization of the SSE and the average silhouette width values (Fig. S2).

The identified clusters were characterized by nearly unique sets of enriched GO terms and reflected the identified phases of the gametogenesis process well (Fig. 2). Cluster 1 grouped genes expressed in the vegetative phase with decreasing expression levels during gametogenesis. Cluster 2 was the only cluster with high expression during the determination phase but low expression during the differentiation and swarming phase. Cluster 3 and 4 grouped genes with high expression during the differentiation phase. The main difference between both clusters consisted of genes of cluster 3 being downregulated during the swarming phase. In contrast, genes in clusters 4 were characterized by a high expression level during the swarming phase also. Cluster 5 contains the least number of genes, upregulated only in the swarming phase.

**Figure 2.**
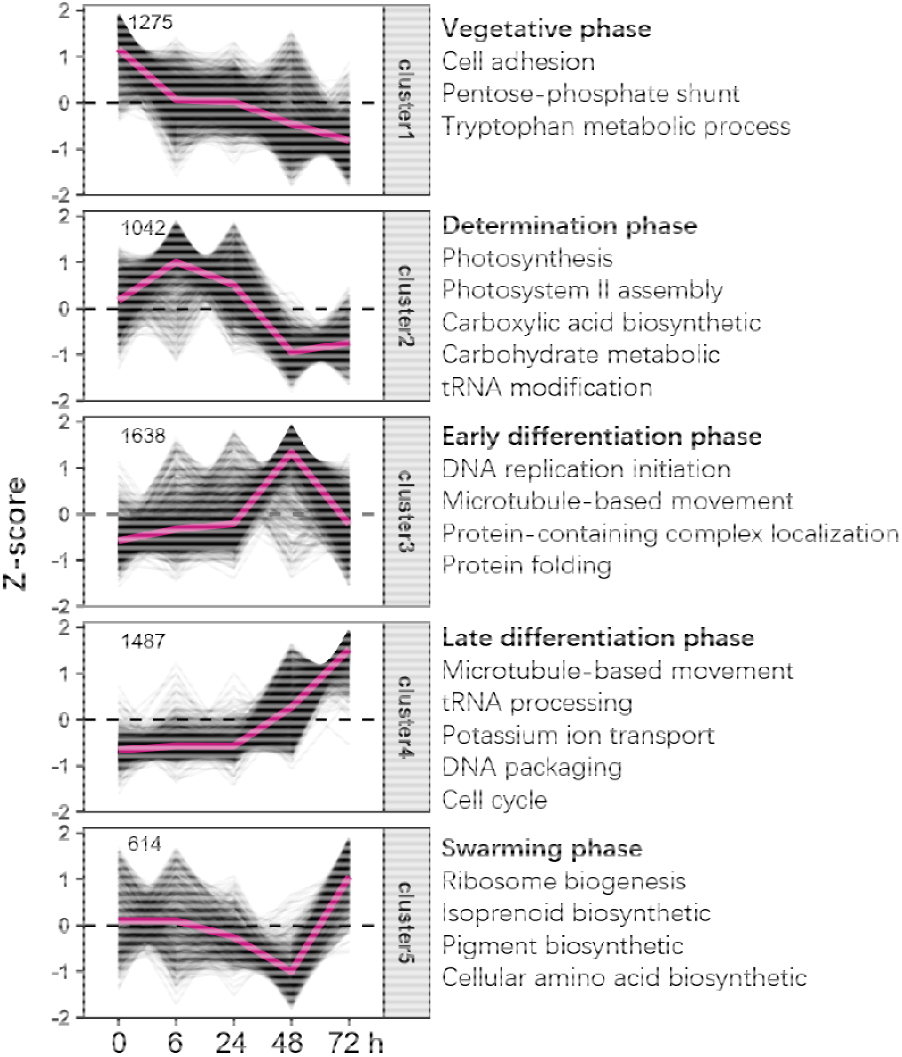
Clusters of differentially expressed genes (DEG) identified during the different phases of gametogenesis at 0, 6, 24, 48 and 72h. Clusters are presented according their maximal expression in the different phases of gametogenesis. A selection of the most relevant overrepresented GO terms (p<0.01 and summarized by REVIGO) is presented for each cluster. We refer to Table S5 for the full list enriched GO terms.

DEGs related to photosynthesis and carboxylic acid biosynthesis were upregulated during the determination phase. Next, genes involved in DNA replication and microtubule-based movements have a high expression during the early differentiation process of gametogenesis as well as during the gamete formation. However, the different enriched GO terms between the two clusters apply primarily to DNA replication which indicates that DNA replication is completed in the early phase of the differentiation process, before gametes formation starts. And the final swarming phase was mainly relevant to the pigment synthesis and cellular amino acid biosynthesis which was the final step for the gametes formation.

### Initiation of gametogenesis versus response to fragmentation

By adding an additional control group whereby thalli were fragmented but the sporulation inhibitors were not removed (CNW, chopping no washing), we aimed to disentangle the effect of fragmentation on gene expression from the induction of gametogenesis. By analyzing DEGs between CW and CNW group at 6 h and 24 h, we identified 901 and 1137 significantly differential expressed genes (FDR<0.05, log2FC>1 or <-1) at 6 h and 24 h, respectively. DEGs included 493 up-regulated and 408 down-regulated genes at 6 h and 652 up-regulated and 485 down-regulated genes at 24 h. Nearly half of the DEGs at 6 h have the same expression profile at 24 h compared to the CNW group (Fig. 3). Of the 493 up-regulated and 408 down-regulated genes at 6 h, 206 and 219 genes showed the same expression profiles at 24 h, indicating that DEGs with a function in the determination phase tended to have continuous expression profiles during early gametogenesis. The enriched GO terms for the up-regulated DEGs at 6 h are mainly related to the photosynthesis, ATP biosynthetic process and oxidation-reduction processes (Fig. 4), which are in accordance with cluster 2 enrichment. In contrast, GO terms significantly enriched at 24 h are mainly related to the microtubule-based movement, DNA replication and cilium organization, suggesting that the DNA replication initiation for the gametogenesis starts from 24 h, consistent with the expression pattern of cluster 3 (Fig. 2). The over-represented GO terms for the down-regulated genes were related to the response to oxidative stress and oxidation-reduction process both at 6 h and 24 h after induction of gametogenesis.

**Figure 3.**
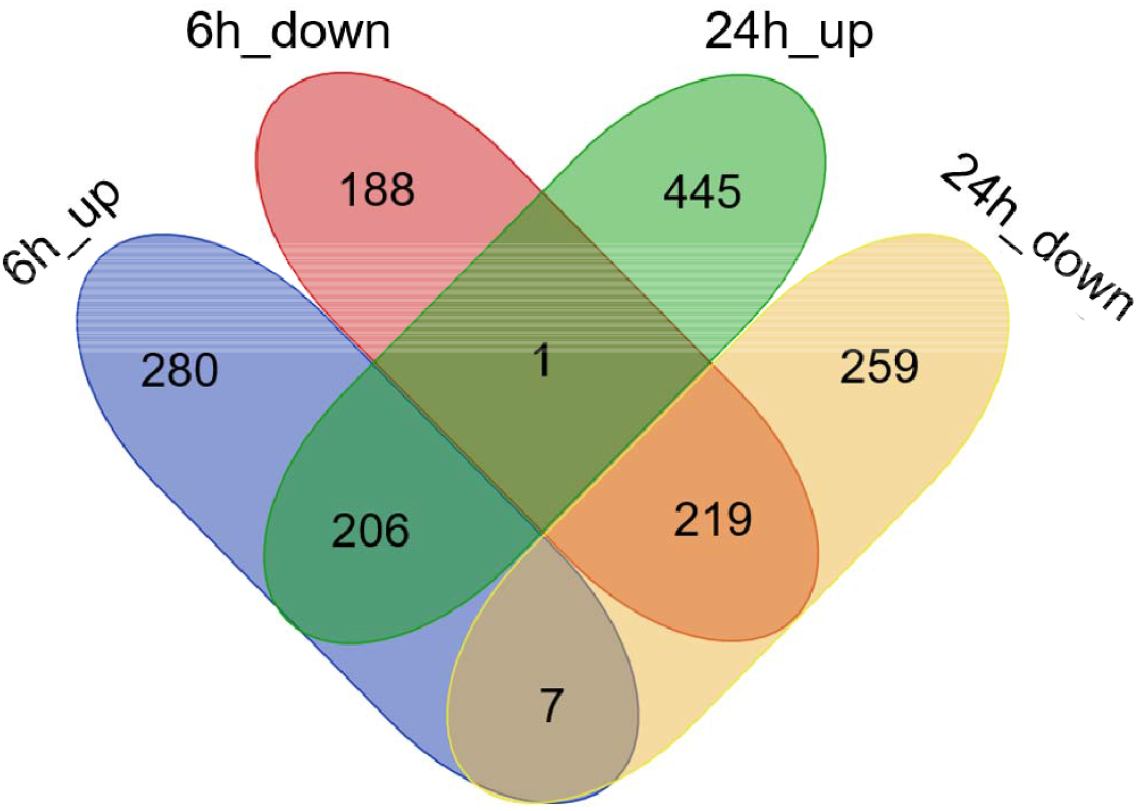
Differentially expressed genes (DEGs) between fully induced thalli (CW) and thalli with the sporulation inhibitors still present (CNW). The Venn diagram depicts the number of up- and downregulated DEGs at 6 h and 24 h.

**Figure 4.**
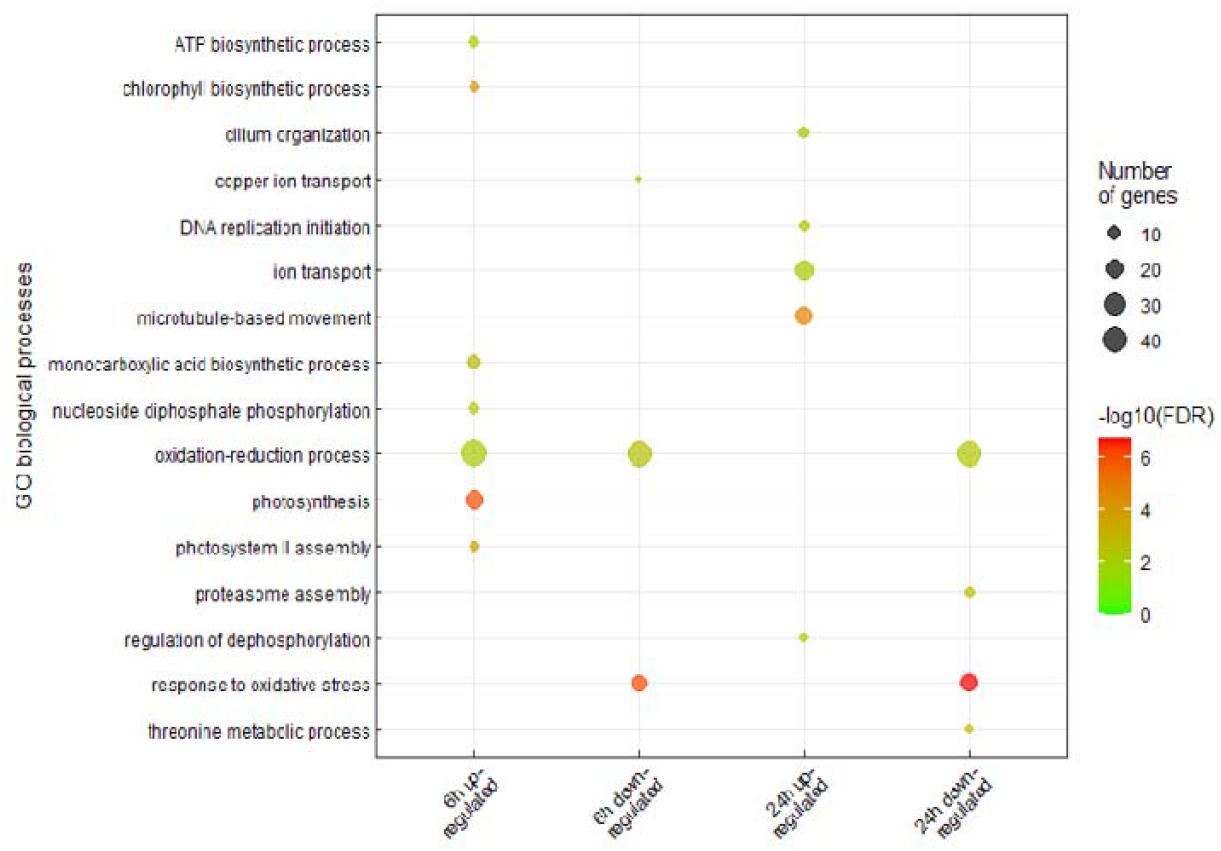
Enriched GO terms associated with DEGs between fully induced thalli (CW) and thalli with the sporulation inhibitors still present (CNW) at 6 h and 24 h respectively.

To identify the possible key initiator for *Ulva* gametogenesis, we analyzed the differentially expressed TFs at 6 h which signifies the early responder to the washing treatment exclusively by comparing the chopping and washing (CW) group with the chopping without washing (CNW) group. We identified 12 up-regulated and 6 down-regulated transcription factors (TFs) at 6 h (Table 1). Several homologs of these TFs, as represented by the top blast hits in the *Ulva*-PLAZA database, are involved in gamete formation in *Arabidopsis* or *Chlamydomonas*.

**Table 1.**
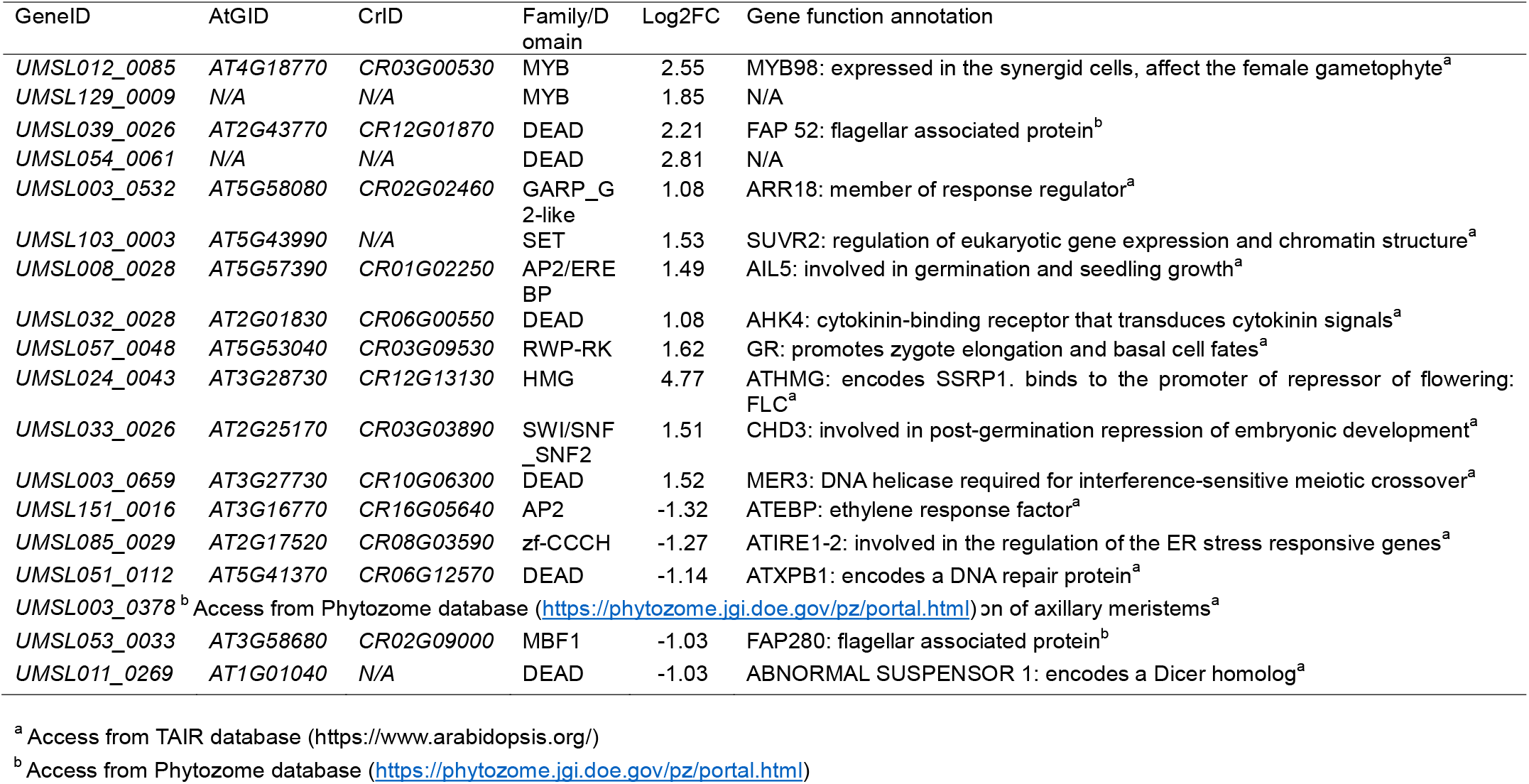
Differential expressed TFs of induced (CW) versus non-induced (CNW) thalli at 6 h. Log2FC>1: upregulated, LogFC<-1: downregulated. The homologs of the TFs of *A. thaliana* and *C. reinhardtii* are listed in the “AtGID” and “CrID” columns respectively. Gene function annotation are summarized from TAIR or Phytozome databases.

| GeneID | AtGID | CrID | Family/D<br>omain | Log2FC | Gene function annotation |
| --- | --- | --- | --- | --- | --- |
| UMSL012_0085 | AT4G18770 | CR03G00530 | MYB | 2.55 | MYB98: expressed in the synergid cells, affect the female gametophyte<sup>a</sup> |
| UMSL129_0009 | N/A | N/A | MYB | 1.85 | N/A |
| UMSL039_0026 | AT2G43770 | CR12G01870 | DEAD | 2.21 | FAP 52: flagellar associated protein<sup>b</sup> |
| UMSL054_0061 | N/A | N/A | DEAD | 2.81 | N/A |
| UMSL003_0532 | AT5G58080 | CR02G02460 | GARP_G<br>2-like | 1.08 | ARR18: member of response regulator<sup>a</sup> |
| UMSL103_0003 | AT5G43990 | N/A | SET | 1.53 | SUVR2: regulation of eukaryotic gene expression and chromatin structure<sup>a</sup> |
| UMSL008_0028 | AT5G57390 | CR01G02250 | AP2/ERE<br>BP | 1.49 | AIL5: involved in germination and seedling growth<sup>a</sup> |
| UMSL032_0028 | AT2G01830 | CR06G00550 | DEAD | 1.08 | AHK4: cytokinin-binding receptor that transduces cytokinin signals<sup>a</sup> |
| UMSL057_0048 | AT5G53040 | CR03G09530 | RWP-RK | 1.62 | GR: promotes zygote elongation and basal cell fates<sup>a</sup> |
| UMSL024_0043 | AT3G28730 | CR12G13130 | HMG | 4.77 | ATHMG: encodes SSRP1. binds to the promoter of repressor of flowering:<br>FLC<sup>a</sup> |
| UMSL033_0026 | AT2G25170 | CR03G03890 | SWI/SNF<br>_SNF2 | 1.51 | CHD3: involved in post-germination repression of embryonic development<sup>a</sup> |
| UMSL003_0659 | AT3G27730 | CR10G06300 | DEAD | 1.52 | MER3: DNA helicase required for interference-sensitive meiotic crossover<sup>a</sup> |
| UMSL151_0016 | AT3G16770 | CR16G05640 | AP2 | -1.32 | ATEBP: ethylene response factor<sup>a</sup> |
| UMSL085_0029 | AT2G17520 | CR08G03590 | zf-CCCH | -1.27 | ATIRE1-2: involved in the regulation of the ER stress responsive genes<sup>a</sup> |
| UMSL051_0112 | AT5G41370 | CR06G12570 | DEAD | -1.14 | ATXPB1: encodes a DNA repair protein<sup>a</sup> |
| UMSL003_0378 | AT1G19270 | CR10G01560 | DEAD | -1.11 | DA1: controls the initiation of axillary meristems<sup>a</sup> |
| UMSL053_0033 | AT3G58680 | CR02G09000 | MBF1 | -1.03 | FAP280: flagellar associated protein<sup>b</sup> |
| UMSL011_0269 | AT1G01040 | N/A | DEAD | -1.03 | ABNORMAL SUSPENSOR 1: encodes a Dicer homolog<sup>a</sup> |

<sup>a</sup> Access from TAIR database (https://www.arabidopsis.org/)
<sup>b</sup> Access from Phytozome database (https://phytozome.jgi.doe.gov/pz/portal.html)

We acquired the homologs of the *Arabidopsis* and *Chlamydomonas* of the upregulated TFs at 6 h by blast. With reported function of the homologs, we tried to find the conserved TFs function in plants gametogenesis. Particularly, an extensive studied TF containing RWP-RK domain aroused our interests for further characterization. The RWP-RK gene in *Chlamydomonas*, which is minus dominance (MID) is responsible for switching on the minus-programme and switching off the plus-programme in gamete differentiation (Lin and Goodenough, 2007). Studies in *Arabidopsis* indicate that the RWP-RK TFs control cell differentiation during female gametophyte development (Tedeschi *et al.*, 2017). We investigated the expression pattern of the RWP-RK TF (*UMSL057_0048*) in more detail by means of qRT-PCR analysis, sampling tissue fragments undergoing gametogenesis every 2 hours. At each timepoints the expression was compared against a control treatment not undergoing gametogenesis. *UMSL057_0048* is up-regulated in the early determination phase and throughout the determination and differentiation phase with maximum expression levels at 6 h (Fig. 5a). More specifically, *UMSL057_0048* shows a quick response to the chopping and washing treatment with slight upregulation at 2 h after induction and significantly upregulation at 4 h.

**Figure 5.**
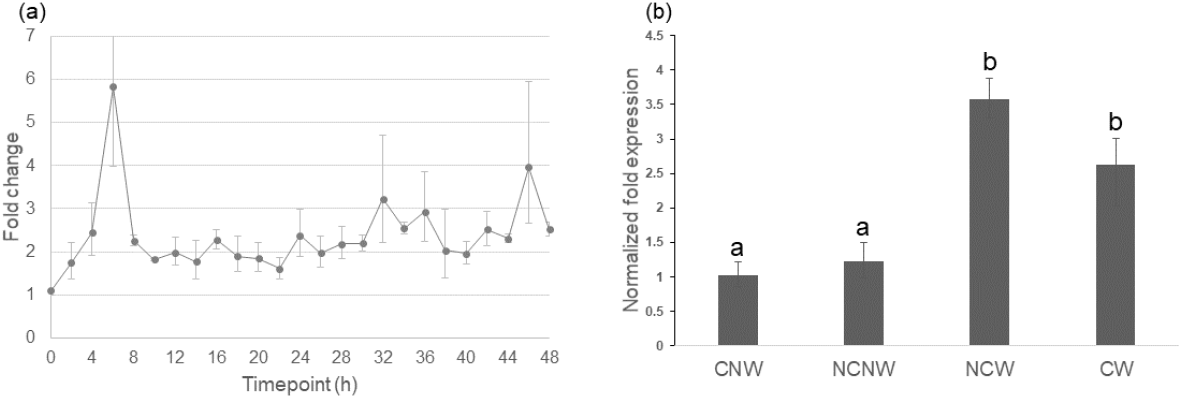
Expression of RWP-RK transcription factor *UMSL057_0048* during gametogenesis. (a). RT-qPCR analysis of *UMSL057_0048* expression level compared to the control group (NCNW) in the first 48 h of gametogenesis. Three biological replicates were used for analysis and the gene expression level was calculated using the standard ddCt method. (b). Expression level of *UMSL057_0048* at 6 h in induced (CW and NCW) versus not induced (CNW and NCNW) thalli. Three biological replicates were used for the experiment and standard ddCt method was applied for analysis. Significant differences are indicated with lowercase (ANOVA test, *P* < 0.05). Stand errors are indicated with errors bars.

In an additional experiment we contrasted the expression at 6 h of *UMSL057_0048* to fragmented thalli from which the SI were not removed (CNW) and unfragmented thalli that were washed to remove the SI in the medium and hence were partially induced (NCW). qRT-PCR results indicate *UMSL057_0048* is not upregulated at 6 h in treatments where gametogenesis is not induced (CNW and NCNW). In contrast in both treatments where gametogenesis is induced (CW and NCW), *UMSL057_0048* is significantly upregulated (Fig. 5b). The *Ulva* genome contains one additional gene with an RWP-RK domain, *UMSL048_0014.* The latter, however, is not expressed (CPM value ~0) during gametogenesis.

## Discussion

The transition from a vegetative to a reproductive state is a key step in the life cycle of *Ulva.* Analysis of the molecular mechanisms underpinning this process is pivotal for efficient seeding technologies and putative genetic improvement for *Ulva* as a commercial crop species. In this study, RNA-seq was applied to analyze the dynamic changes of gene expression of *Ulva mutabilis* and to identify key TFs involved in the initiation of *Ulva* gametogenesis. Transcriptional analysis showed that nearly half (45.4%) of the protein-coding genes predicted in the draft genome are differential expressed between at least 2 timepoints during gametogenesis. The large reprogramming of the transcriptome likely reflects the full transformation of a vegetative cell into 16 gametes (Wichard and Oertel, 2010).

### Dynamic changes in gene regulation during gametogenesis

GO enrichment analysis of DEGs clusters sheds light on the most enriched up- or down-regulated biological process during each of the three phases that characterize gametogenesis in *Ulva.* Cluster 1 contains DEGs that are downregulated during the determination phase and genes in this cluster are enriched in the cell adhesion and pentose-phosphate shunt activity. For cluster 2 which is upregulated during the determination phase, there is a high expression of genes that are associated with photosynthesis, carboxylic acid biosynthesis and ribose phosphate metabolism. This correlates with significant increases in maltose during gametogenesis of *U. prolifera* at 12 h and 24 h is, which are approximately 12-fold change up-regulated compared to the control group (He *et al.*, 2019). Similarly, the upregulated photosynthesis related genes at 24 h and 48 h may cause the increase in chlorophyll content subsequent to the divisions of the chloroplast as demonstrated for *Ulva pertusa* during sporulation (Park, 2020). All clusters represent a marked changes of gene expression in the transition from determination phase (24 h) to differentiation phase (48 h). 36 h after induction is regarded an important checkpoint during gametogenesis after which blade cells are irreversibly committed to gametangium differentiation (Kessler *et al.*, 2017; Stratmann *et al.*, 1996; Wichard and Oertel, 2010). Genes related to DNA replication and microtubule-based movement were significantly up-regulated after this irreversible timepoint. Electron microscope observations indeed demonstrate drastic rearrangements in microtubule morphology between 36 h and 48 h after induction. The microtubule cytoskeleton of somatic cells consists of parallel microtubule bundles arranged mainly in the cortical cytoplasm parallel to the plasmalemma. However, 36 h after induction, the microtubule bundles traverse the cortical cytoplasm converging on a particular area and then the microtubules starting from a pointed site at basal part of gametangium and forming a basket-like configuration with a circular opening at the top after 48 h (Katsaros *et al.*, 2017). Stratmann et al. (1996) analyzed the DNA synthesis during gametogenesis and concluded that DNA replication starts from 25 hours after induction which is consistent with our RNA-seq data. The enriched GO terms of microtubule-based movement in cluster 3 and 4 are the molecular basis of the cytological observation of the cytoskeleton organization. Genes in cluster 5 are upregulated only during the swarming phase and the enriched GO terms mainly involve the synthesis of amino acid and isoprenoid pigment biosynthesis. Isoprenoid pigments biosynthetic pathway catalyzes the synthesis of essential pigments of the photosystem structure in plant cells (Couso *et al.*, 2012). We speculate that the upregulation of isoprenoid biosynthetic pathway is related to the reconstruction of the photosynthesis system for the gamete formation.

### Transcription factors associated with gametogenesis

Transcription factors play a pivotal role in the gene expression networks. TFs were surveyed in the developmental program of gametogenesis that involves extensive cellular morphogenesis and subsequent cell division and proliferation. *Ulva* gametogenesis was initiated by fragmentation and subsequent removal of SI-1 and SI-2. As shown by Stratmann et al. (1996), fragmentation of growing gametophytes itself does not initiate gametogenesis and the cells remain in a “vegetative” state after 72 h because of the rapidly excreted SI-1 in the culture medium suppresses gametogenesis during the determination phase. To remove the effect of fragmentation and to identify a putative regulator initiating gametogenesis in *Ulva*, we designed an experiment where thalli are fragmented but the sporulation inhibitors are not removed. With this control group, we identified 12 up-regulated and 6 down-regulated TFs at 6 h after induction of gametogenesis which constitute about 4-5% of the TF repertoire of *U. mutabilis.*

We selected the RWP-RK TF (*UMSL057_0048*) for further analysis because studies indicate a conserved role of RWP-RK domain containing TFs in gametogenesis process (Chardin *et al.*, 2014; Rovekamp *et al.*, 2016; Tedeschi *et al.*, 2017; Yin *et al.*, 2020). Nevertheless, the other TFs identified in this study might be potentially interesting as well. For example, Myb domain protein 98 (MYB98, *UMSL012_0085*) is involved in the regulation of synergid differentiation in angiosperms (Kasahara *et al.*, 2005; Punwani *et al.*, 2007). Similarly, homologs of high nobility group transcription factor (*UMSL024_0043*) are relevant for the transition of a vegetative to a reproductive phase (Klepikova *et al.*, 2015; Pfab *et al.*, 2018; Searle *et al.*, 2006). The homolog of *UMSL024_0043*, which is the most up-regulated gene compared to the control group at 6 h, encodes SSRP1, which is component of the facilitates chromatin transcription (FACT) complex in Arabidopsis. FACT is a conserved heterodimeric histone chaperone among eukaryotes and facilitates expression of *FLC* (FLOWERING LOCUS C) by binding to the promoter of *FLC*, which adjusts the switch from vegetative to reproductive development in Arabidopsis (*Grasser, 2020; Lolas et al., 2010*). Reduced amounts of SSRP1 result in decreased expression of *FLC*, thus accelerating flowering (Lolas *et al.*, 2010). In contrast, *UMSL024_0043* is up-regulated during gametogenesis and we did not identify a homolog of *FLC* in the *Ulva* genome. Thus, the specific function and the target genes of *UMSL024_0043* will need to be confirmed experimentally.

One particular TF (encoded by *UMSL057_0048*) containing a RWP-RK protein motif that is important in DNA binding, is upregulated at 6h. Remarkably, minus dominance (MID), homolog of RWP-RK TFs in *C. reinhardtii*, has been described to be required for gametogenesis (Ferris *et al.*, 1997). In addition, similar functions of RWP-RK TFs during gametogenesis were reported in many other plants (Tedeschi *et al.*, 2017; Waki *et al.*, 2011). RWP-RK transcription factors are found throughout the Viridiplantae. Phylogenetic analysis divided the RWP-RK homologs into 4 subfamilies, RKD(A), RKD(B), RKD(C) and NLP subfamily (Chardin *et al.*, 2014). Previous studies on algae and higher plants showed that members of NLP subfamily were reported as the early regulators of cellular response to N supply (Camargo *et al.*, 2007; Castaings *et al.*, 2009) and RKD homologs plays evolutionarily conserved roles in germ cell differentiation (Camargo *et al.*, 2007; Castaings *et al.*, 2009; Chardin *et al.*, 2014; Koi *et al.*, 2016; Lin and Goodenough, 2007; Yin *et al.*, 2020). Unlike other organisms which have multiple RWP-RK transcription factors, only two RWP-RK transcription factors are identified in the *U. mutabilis* slender genome. A recent study on the *Ulva partita* mating type locus structure reported three RWP-RK TFs in the *U. partita* genome, including one RWP-RK TF located in the mating type minus locus with weak homology with MID gene (Yamazaki *et al.*, 2017). The genome applied in our analysis is mating type plus, so the gene located in the mating type minus is not characterized. UMSL048_0014 contains a conserved GAF domain that is shared in the NLP subfamily. Two members of this subfamily were identified as nitrogen response TF in *C. reinhardtii* (CreNIT2) and *Volvox carteri* (VcaNIT2) (Camargo *et al.*, 2007; Castaings *et al.*, 2009). Interestingly, the gametogenesis of *C. reinhardtii* is activated by the N-removal induction, while nitrogen concentration seems to have a positive effect on reproduction in *Ulva* (Gao *et al.*, 2017). This may suggest that *UMSL048_0014* may have a function in nitrogen metabolism pathway only. UMSL057*_*0048 identified in our study shares a conserved protein domain with RKD(A) family. In our study, *UMSL057_0048* is exclusively upregulated in the gametogenesis group (CW and NCW) and shows upregulation at 2 h after fragmentation and washing treatments. These findings indicate that RKD family member *UMSL057_0048* plays an essential role in *Ulva* gametogenesis.

Compared to *UMSL057_0048, UMSL048_0014* contains a truncated GAF domain, which is conserved in the NIN-like proteins (NLPs) subfamily and this subfamily is stated to be the early regulators of cellular response to N supply (Camargo *et al.*, 2007; Castaings *et al.*, 2009; Chardin *et al.*, 2014). In contrast, *UMSL057_0048* contains a conserved protein domain, which Chardin et al. referred to as motif 12, which is conserved in RKD subfamily (Fig. 6) and this subfamily is involved in gametogenesis (Camargo *et al.*, 2007; Castaings *et al.*, 2009; Chardin *et al.*, 2014; Koi *et al.*, 2016; Lin and Goodenough, 2007).

**Figure 6.**
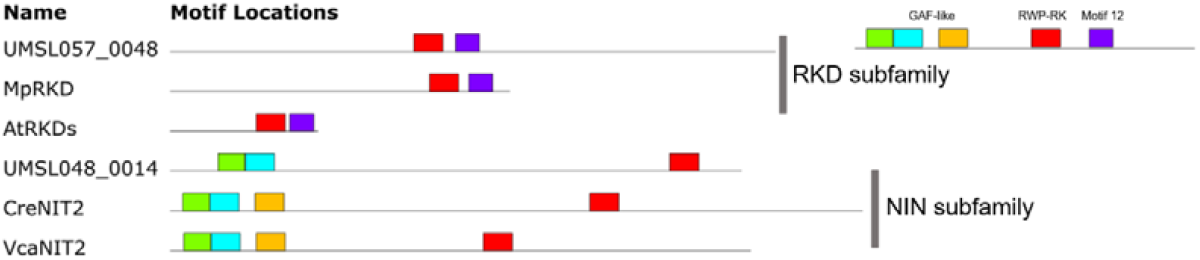
Comparison of domain organization between RWP-RK proteins. “Motif 12” refers to the same motif identified by Chardin et al. (2014). Model organisms in which the RWP-RK protein were functionally validated were chosen as reference. UMSL, *Ulva mutabilis* slender (+); Mp, *Marchantia polymorpha;* At, *Arabidopsis thaliana;* Cre, *Chlamydomonas reinhardtii*; Vca, *Volvox carteri*; Amino acid sequences and accession numbers are listed in Table S6.

## Conclusive remarks

Our present work provides an important resource to study gametogenesis regulatory framework in a model organism from which insights can be translated into commercially important *Ulva* species. Transcriptomic profiling helps us understand the gene regulatory networks during *Ulva* gametogenesis. The CNW group provides useful information to understand *Ulva* response to fragmentation and combined fragmentation and washing factors respectively. The genetic transformation system is available in *U. mutabilis* and further genome editing technology is under development (Oertel *et al.*, 2015), which will allow functional characterisation of gametogenesis-related TFs in future experiments. The understanding of the mechanisms of inducing *Ulva* gametogenesis would be helpful to the commercial cultivation of *Ulva*, extending to other seaweeds. In addition, *Ulva* has developed multicellularity independently from land plants and our study suggests that some TFs play a conserved role in reproduction throughout the green lineage. Therefore, further study of gametogenesis in *Ulva* should provide us with evolutionary insights into control of the sexual reproduction process for the Viridiplantae in general.

## Supporting information

Supplemental Figuires 1 -2

## Supplementary data

Figure S1. PCA analysis for the all the sequenced samples.

Figure S2. Analysis for the optimal number of the clusters for DEGs. (a): The sum of the squared distance between each member of a cluster and its cluster centroid (SSE) analysis. (b): The silhouette value analysis describes how similar a gene is to its own cluster compared to other clusters.

Table S1. Primer sequences used for qRT-PCR analysis.

Table S2. The total number of mapped reads for each sample.

Table S3. The raw reads count for each gene.

Table S4. Pearson R2 test for all the samples.

Table S5. Full enriched GO terms for each cluster.

Table S6. Amino acid sequences and accession numbers for the protein domain analysis.

## Acknowledgements

X.L. is indebted to the China Scholarship Council (201504910698) and Ghent University Special Research Fund (BOF-16/CHN/023) for a PhD grant and to COST Action FA1406 (Phycomorph) for a Short-Term Scientific Mission to Friedrich Schiller University Jena. J.B. thanks the Research Foundation – Flanders (FWO) for a postdoctoral fellowship (12T3418N) and Ghent University Special Research Grant (BOF20/PDO/016). T.W. was funded by the Deutsche Forschungsgemeinschaft through Grant No. SFB 1127/2 ChemBioSys–239748522. This work makes use of resources and facilities provided by UGent as part of the Belgian contribution to EMBRC-ERIC (FWO GOH3817N) and the Betty Moore Foundation for a SYMBIOSIS grant (nr. 4546891618).

## Author contribution

X.L., and O.D.C. designed the study. X.L., T.W. and S.D. performed experiments. X.L., J.B. and K.B. performed the transcriptional analysis. X.L. wrote the manuscript with contributions from all authors.

## Data availability

The raw reads generated in this study are available on SRAXXX.

## Notes

### Competing Interest Statement

The authors have declared no competing interest.

