## Supplemental Figuires 1 -2 for "Gametogenesis in the green seaweed *Ulva mutabilis* coincides with massive transcriptional restructuring"

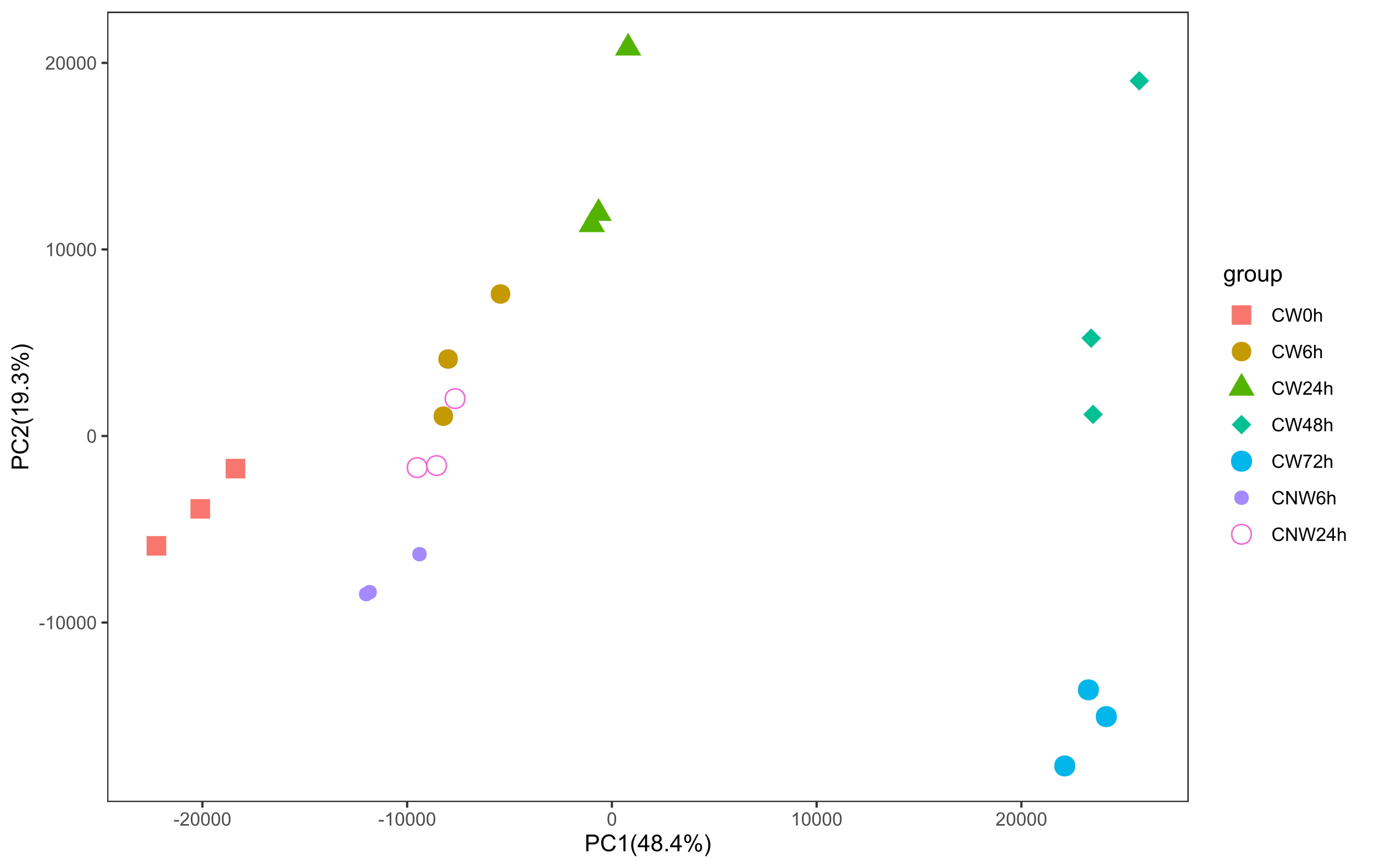

Figure S1. PCA analysis for the all the samples. Three replicates of each group are indicated by same color and shape.

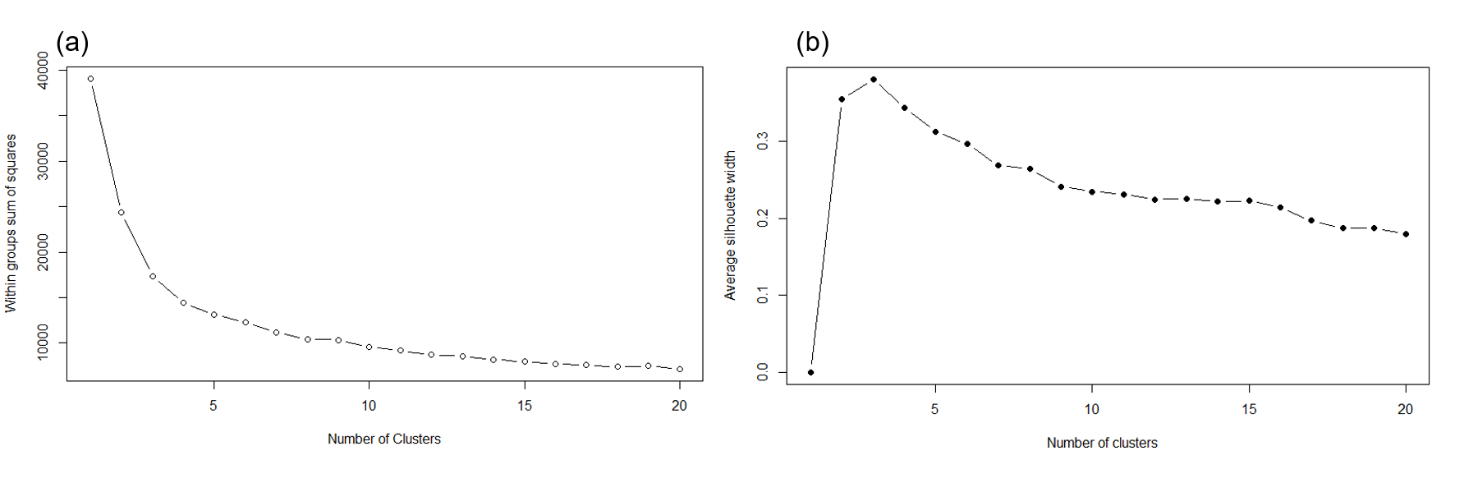

Figure S2. Analysis for the optimal number of the clusters for DEGs. (a): The sum of the squared distance between each member of a cluster and its cluster centroid (SSE) analysis. (b): The silhouette value analysis describes how similar a gene is to its own cluster compared to other clusters.
